## Supplemental Materials for "Gliding towards an understanding of the origin of flight in bats" (from another version of this preprint: version 3, 2023-10-06)

Supplementary Information for
**Gliding towards an understanding of the origin of flight in bats**

This PDF file includes:

**Supplementary Results**

Figure S1: PCA with fossil and extant species of interest labeled.

Table S1: Loadings for forelimb and hind limb PCAs with extant mammals and extinct bats.

Table S2: Statistical results for OUM model fitting analyses on the extant-only dataset.

Table S3: Optima and bootstrapping results for best-supported OUM model.

Table S4: Statistical results for OUM model fitting analyses on the extant+extinct dataset.

Table S5: Optima values for the best-fitting OUM model results for extant+extinct dataset.

Table S6: Fits of multivariate evolutionary models to extant+extinct dataset.

Table S7: Summary of the best-fitting univariate OUwie models with interpretation of the optima in the context of our initial predictions.

**Other supplementary materials for this manuscript**

Data File S1: Morphological dataset for 231 mammal species and 4 extinct bats (.csv file).

**Figure S1: PCA with fossil and extant species of interest labeled.** Morphospaces consisting of PCs 1 and 2 of our extant and extinct species' forelimb dataset for a) forelimbs and b) hind limbs with arrows pointing to the dermopteran species *Galeopteras variegatus* and *Cynocephalus volans*, the common vampire bat *Desmodus rotundus*, and the oldest fossil bat, *Onchonycteris finneyi*.

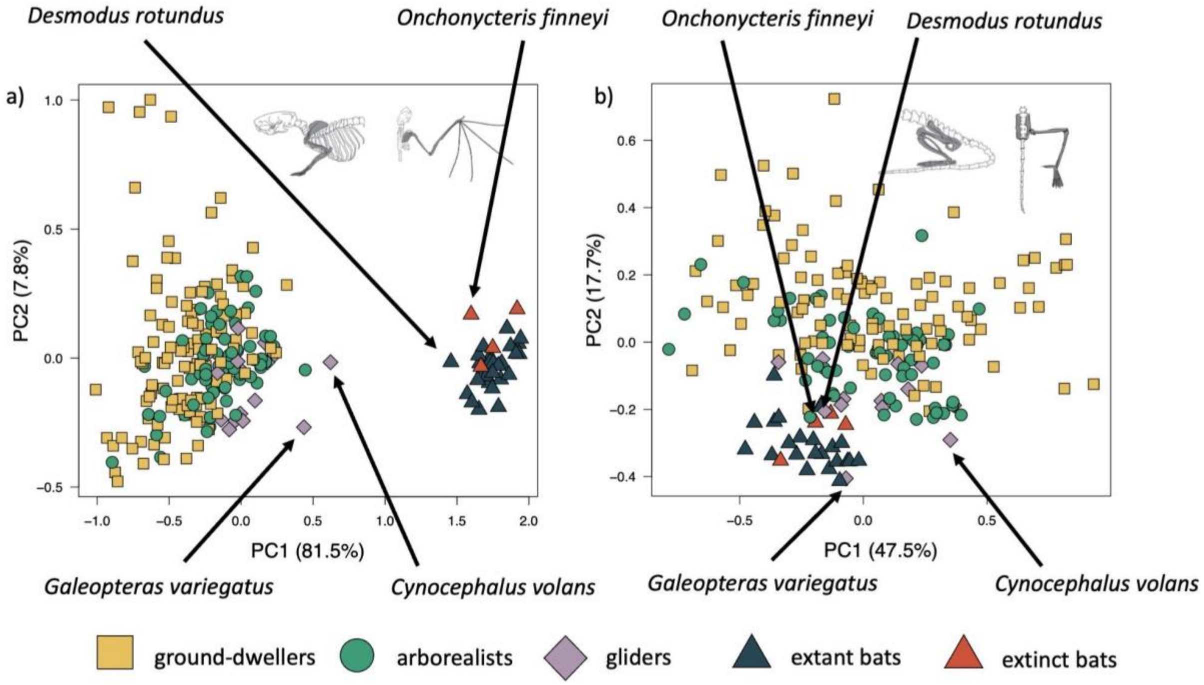

**Table S1: Loadings for forelimb and hind limb PCAs with extant mammals and extinct bats.** Principal component analysis loadings for our a) 14 forelimb traits and b) 10 hind limb traits (pes were excluded due to insufficient data). The five most extreme loadings are bolded for PC1 and PC2.

| | **component loadings** | | |
|---|---|---|---|
| **a) forelimb traits** | **PC1** | **PC2** | **PC3** |
| scapula length (sl) | 0.056 | 0.186 | 0.079 |
| scapula height (sh) | 0.099 | 0.174 | **0.258** |
| humerus length (hl) | **0.212** | 0.046 | -0.076 |
| humerus shaft width (hsw) | 0.088 | **0.342** | -0.186 |
| humerus proximal width (hpw) | 0.085 | 0.259 | 0.000 |
| humerus distal width (hdw) | 0.045 | **0.344** | **-0.273** |
| ulna length (ul) | 0.256 | 0.021 | -0.089 |
| ulna olecranon length (uol) | -0.066 | **0.394** | **0.329** |
| radius length (rl) | **0.300** | -0.064 | -0.115 |
| 3rd metacarpal length (mcl) | **0.518** | -0.135 | **0.703** |
| 3rd metacarpal width (mcw) | 0.068 | **0.452** | 0.142 |
| 3rd proximal phalanx length (ppl) | **0.447** | -0.118 | -0.183 |
| 3rd proximal phalanx width (ppw) | 0.004 | **0.483** | -0.017 |
| 3rd intermediate phalanx length (ipl) | **0.540** | 0.044 | **-0.367** |
| **proportion explained (%)** | 81.5 | 7.8 | 2.9 |

| | **component loadings** | | |
|---|---|---|---|
| **b) hind limb traits** | **PC1** | **PC2** | **PC3** |
| pelvis length (pel) | 0.188 | 0.189 | 0.262 |
| ilium length (il) | 0.151 | 0.083 | **0.341** |

| | | | |
|---|---|---|---|
| ischium length (isl) | 0.250 | **0.357** | 0.180 |
| femur length (fl) | 0.171 | -0.25 | **0.331** |
| femur shaft width (fsw) | 0.131 | 0.184 | 0.182 |
| femur distal width (fdw) | 0.141 | 0.164 | 0.124 |
| tibia length (tl) | 0.272 | -0.257 | 0.239 |
| tibia proximal width (tpw) | 0.151 | 0.141 | 0.091 |
| tibia midshaft width (tmw) | 0.235 | 0.134 | 0.118 |
| fibula length (fbl) | **0.301** | -0.204 | **0.299** |
| 3rd metatarsal length (mtl) | **0.475** | 0.088 | -0.211 |
| 3rd metatarsal width (mtw) | **0.335** | **0.282** | **-0.448** |
| 3rd pedal proximal phalanx length (ppxl) | **0.310** | **-0.378** | -0.074 |
| 3rd pedal proximal phalanx width (ppxw) | 0.218 | **0.295** | -0.273 |
| 3rd intermediate pedal phalanx length (ipxl) | **0.297** | **-0.494** | **-0.362** |
| **proportion explained (%)** | 47.5 | 17.8 | 11.3 |

**Table S2: Statistical results for OUM model fitting analyses on the extant-only dataset.** Comparisons of the best-fitting OUM models (bolded) using AICc scores to assess fit and AICcW weights to asses relative model support for a) 14 forelimb traits and b) 15 hind limb traits. All models were fit using functions in the *OUwie R* package. Trait abbreviations are explained in table S1.

### a. forelimb

| sl | fit | delta | w |
|---|---|---|---|
| BM1 | -382.99 | 134.24 | 0.00 |
| OU1 | -501.85 | 15.39 | 0.00 |
| $OUM_{loc3a}$ | -510.78 | 6.45 | 0.02 |
| **$OUM_{loc3b}$** | **-517.24** | **0.00** | **0.49** |
| $OUM_{loc3c}$ | -516.17 | 1.07 | 0.28 |
| $OUM_{loc4}$ | -515.57 | 1.67 | 0.21 |

| sh | fit | delta | w |
|---|---|---|---|
| BM1 | -304.60 | 92.15 | 0.00 |
| OU1 | -368.87 | 27.88 | 0.00 |
| $OUM_{loc3a}$ | -376.88 | 19.87 | 0.00 |
| $OUM_{loc3b}$ | -394.81 | 1.95 | 0.19 |
| $OUM_{loc3c}$ | -395.61 | 1.14 | 0.29 |
| **$OUM_{loc4}$** | **-396.75** | **0.00** | **0.51** |

| hl | fit | delta | w |
|---|---|---|---|
| BM1 | -418.82 | 89.49 | 0.00 |
| OU1 | -430.16 | 78.15 | 0.00 |
| $OUM_{loc3a}$ | -503.40 | 4.91 | 0.08 |
| $OUM_{loc3b}$ | -493.66 | 14.66 | 0.00 |
| $OUM_{loc3c}$ | -491.74 | 16.57 | 0.00 |
| **$OUM_{loc4}$** | **-508.31** | **0.00** | **0.92** |

| hdw | fit | delta | w |
|---|---|---|---|
| BM1 | -395.19 | 43.43 | 0.00 |
| OU1 | -435.24 | 3.38 | 0.07 |
| $OUM_{loc3a}$ | -438.61 | 0.01 | 0.36 |
| **$OUM_{loc3b}$** | **-438.62** | **0.00** | **0.36** |
| $OUM_{loc3c}$ | -434.91 | 3.71 | 0.06 |
| $OUM_{loc4}$ | -436.91 | 1.71 | 0.15 |

| hsw | fit | delta | w |
|---|---|---|---|
| BM1 | -350.21 | 82.77 | 0.00 |
| OU1 | -410.29 | 22.69 | 0.00 |
| $OUM_{loc3a}$ | -426.30 | 6.69 | 0.02 |
| **$OUM_{loc3b}$** | **-432.98** | **0.00** | **0.65** |
| $OUM_{loc3c}$ | -424.81 | 8.17 | 0.01 |
| $OUM_{loc4}$ | -431.55 | 1.44 | 0.32 |

| hpw | fit | delta | w |
|---|---|---|---|
| BM1 | -451.41 | 94.80 | 0.00 |
| OU1 | -511.63 | 34.59 | 0.00 |
| $OUM_{loc3a}$ | -527.98 | 18.23 | 0.00 |
| **$OUM_{loc3b}$** | **-546.21** | **0.00** | **0.40** |
| $OUM_{loc3c}$ | -545.50 | 0.72 | 0.28 |
| $OUM_{loc4}$ | -545.77 | 0.44 | 0.32 |

| rl | fit | delta | w |
|---|---|---|---|
| BM1 | -369.45 | 85.45 | 0.00 |
| OU1 | -368.87 | 86.03 | 0.00 |
| $OUM_{loc3a}$ | -442.25 | 12.66 | 0.00 |
| $OUM_{loc3b}$ | -437.74 | 17.17 | 0.00 |
| $OUM_{loc3c}$ | -441.09 | 13.81 | 0.00 |
| **$OUM_{loc4}$** | **-454.91** | **0.00** | **1.00** |

| ul | fit | delta | w |
|---|---|---|---|
| BM1 | -410.89 | 113.68 | 0.00 |
| OU1 | -414.76 | 109.80 | 0.00 |
| $OUM_{loc3a}$ | -498.02 | 26.54 | 0.00 |
| $OUM_{loc3b}$ | -506.08 | 18.49 | 0.00 |
| $OUM_{loc3c}$ | -509.29 | 15.28 | 0.00 |
| **$OUM_{loc4}$** | **-524.57** | **0.00** | **1.00** |

| uol | fit | delta | w |
|---|---|---|---|
| BM1 | -272.40 | 73.47 | 0.00 |
| OU1 | -317.22 | 28.65 | 0.00 |
| **$OUM_{loc3a}$** | **-345.87** | **0.00** | **0.58** |
| $OUM_{loc3b}$ | -342.30 | 3.57 | 0.10 |
| $OUM_{loc3c}$ | -331.90 | 13.97 | 0.00 |
| $OUM_{loc4}$ | -344.71 | 1.16 | 0.32 |

| mcl | fit | delta | w |
|---|---|---|---|
| BM1 | -307.39 | 55.35 | 0.00 |

| mcw | fit | delta | w |
|---|---|---|---|
| BM1 | -313.12 | 40.58 | 0.00 |

| ppl | fit | delta | w |
|---|---|---|---|
| BM1 | -308.03 | 81.41 | 0.00 |

| | | | |
|---|---|---|---|
| OU1 | -305.34 | 57.40 | 0.00 |
| OUM$_{loc3a}$ | -333.09 | 29.64 | 0.00 |
| OUM$_{loc3b}$ | -359.99 | 2.75 | 0.16 |
| **OUM$_{loc3c}$** | **-362.74** | **0.00** | **0.63** |
| OUM$_{loc4}$ | -360.58 | 2.16 | 0.21 |

| | | | |
|---|---|---|---|
| OU1 | -351.03 | 2.67 | 0.11 |
| OUM$_{loc3a}$ | -348.07 | 5.64 | 0.03 |
| **OUM$_{loc3b}$** | **-353.70** | **0.00** | **0.44** |
| OUM$_{loc3c}$ | -352.06 | 1.65 | 0.19 |
| OUM$_{loc4}$ | -352.42 | 1.29 | 0.23 |

| | | | |
|---|---|---|---|
| OU1 | -306.13 | 83.31 | 0.00 |
| OUM$_{loc3a}$ | -366.98 | 22.45 | 0.00 |
| OUM$_{loc3b}$ | -386.09 | 3.35 | 0.16 |
| OUM$_{loc3c}$ | -367.24 | 22.20 | 0.00 |
| **OUM$_{loc4}$** | **-389.44** | **0.00** | **0.84** |

| **ppw** | **fit** | **delta** | **w** |
|---|---|---|---|
| BM1 | -290.73 | 53.24 | 0.00 |
| **OU1** | **-343.97** | **0.00** | **0.51** |
| OUM$_{loc3a}$ | -340.07 | 3.90 | 0.07 |
| OUM$_{loc3b}$ | -342.91 | 1.05 | 0.30 |
| OUM$_{loc3c}$ | -339.40 | 4.56 | 0.05 |
| OUM$_{loc4}$ | -339.78 | 4.18 | 0.06 |

| **ipl** | **fit** | **delta** | **w** |
|---|---|---|---|
| BM1 | -221.03 | 111.43 | 0.00 |
| OU1 | -219.46 | 113.00 | 0.00 |
| OUM$_{loc3a}$ | -296.07 | 36.39 | 0.00 |
| OUM$_{loc3b}$ | -323.22 | 9.24 | 0.01 |
| OUM$_{loc3c}$ | -317.39 | 15.07 | 0.00 |
| **OUM$_{loc4}$** | **-332.46** | **0.00** | **0.99** |

### b. hind limb

| **fbl** | **fit** | **delta** | **w** |
|---|---|---|---|
| BM1 | -314.06 | 34.02 | 0.00 |
| OU1 | -341.05 | 7.03 | 0.02 |
| OUM$_{loc3a}$ | -340.55 | 7.53 | 0.01 |
| OUM$_{loc3b}$ | -344.93 | 3.15 | 0.13 |
| OUM$_{loc3c}$ | -346.23 | 1.86 | 0.24 |
| **OUM$_{loc4}$** | **-348.08** | **0.00** | **0.60** |

| **fdw** | **fit** | **delta** | **w** |
|---|---|---|---|
| BM1 | -459.10 | 93.67 | 0.00 |
| OU1 | -533.90 | 18.87 | 0.00 |
| OUM$_{loc3a}$ | -549.84 | 2.93 | 0.10 |
| OUM$_{loc3b}$ | -551.60 | 1.17 | 0.23 |
| **OUM$_{loc3c}$** | **-552.77** | **0.00** | **0.42** |
| OUM$_{loc4}$ | -551.80 | 0.97 | 0.26 |

| **fl** | **fit** | **delta** | **w** |
|---|---|---|---|
| BM1 | -390.02 | 37.29 | 0.00 |
| OU1 | -408.25 | 19.06 | 0.00 |
| OUM$_{loc3a}$ | -425.56 | 1.75 | 0.28 |
| OUM$_{loc3b}$ | -421.54 | 5.77 | 0.04 |
| OUM$_{loc3c}$ | -416.18 | 11.13 | 0.00 |
| **OUM$_{loc4}$** | **-427.31** | **0.00** | **0.68** |

| **fsw** | **fit** | **delta** | **w** |
|---|---|---|---|
| BM1 | -413.39 | 83.94 | 0.00 |
| OU1 | -488.60 | 8.74 | 0.01 |
| **OUM$_{loc3a}$** | **-497.33** | **0.00** | **0.52** |
| OUM$_{loc3b}$ | -492.90 | 4.43 | 0.06 |
| OUM$_{loc3c}$ | -495.90 | 1.44 | 0.26 |

| **il** | **fit** | **delta** | **w** |
|---|---|---|---|
| BM1 | -425.75 | 51.39 | 0.00 |
| OU1 | -468.39 | 8.75 | 0.01 |
| OUM$_{loc3a}$ | -471.22 | 5.93 | 0.03 |
| **OUM$_{loc3b}$** | **-477.14** | **0.00** | **0.52** |
| OUM$_{loc3c}$ | -473.59 | 3.55 | 0.09 |

| **ipxl** | **fit** | **delta** | **w** |
|---|---|---|---|
| BM1 | -95.48 | 97.23 | 0.00 |
| OU1 | -182.86 | 9.84 | 0.00 |
| OUM$_{loc3a}$ | -191.41 | 1.30 | 0.27 |
| **OUM$_{loc3b}$** | **-192.70** | **0.00** | **0.51** |
| OUM$_{loc3c}$ | -186.99 | 5.72 | 0.03 |

| | | | |
|---|---|---|---|
| OUM$_{loc4}$ | -494.89 | 2.44 | 0.16 |

| | | | |
|---|---|---|---|
| OUM$_{loc4}$ | -476.37 | 0.77 | 0.36 |

| | | | |
|---|---|---|---|
| OUM$_{loc4}$ | -190.67 | 2.04 | 0.19 |

| **isl** | **fit** | **delta** | **w** |
|---|---|---|---|
| BM1 | -357.04 | 60.19 | 0.00 |
| OU1 | -392.70 | 24.54 | 0.00 |
| OUM$_{loc3a}$ | -412.19 | 5.04 | 0.06 |
| **OUM$_{loc3b}$** | **-417.23** | **0.00** | **0.71** |
| OUM$_{loc3c}$ | -411.63 | 5.60 | 0.04 |
| OUM$_{loc4}$ | -414.57 | 2.67 | 0.19 |

| **mtl** | **fit** | **delta** | **w** |
|---|---|---|---|
| BM1 | -232.90 | 17.42 | 0.00 |
| OU1 | -243.85 | 6.47 | 0.02 |
| OUM$_{loc3a}$ | -249.63 | 0.69 | 0.31 |
| **OUM$_{loc3b}$** | **-250.32** | **0.00** | **0.44** |
| OUM$_{loc3c}$ | -246.06 | 4.26 | 0.05 |
| OUM$_{loc4}$ | -248.60 | 1.72 | 0.18 |

| **mtw** | **fit** | **delta** | **w** |
|---|---|---|---|
| BM1 | -87.72 | 144.14 | 0.00 |
| OU1 | -215.33 | 16.52 | 0.00 |
| **OUM$_{loc3a}$** | **-231.86** | **0.00** | **0.50** |
| OUM$_{loc3b}$ | -230.38 | 1.47 | 0.24 |
| OUM$_{loc3c}$ | -226.54 | 5.31 | 0.04 |
| OUM$_{loc4}$ | -230.20 | 1.65 | 0.22 |

| **pel** | **fit** | **delta** | **w** |
|---|---|---|---|
| BM1 | -437.41 | 62.83 | 0.00 |
| OU1 | -483.10 | 17.14 | 0.00 |
| OUM$_{loc3a}$ | -491.45 | 8.79 | 0.01 |
| OUM$_{loc3b}$ | -499.65 | 0.59 | 0.35 |
| **OUM$_{loc3c}$** | **-500.24** | **0.00** | **0.48** |
| OUM$_{loc4}$ | -498.11 | 2.13 | 0.16 |

| **ppxl** | **fit** | **delta** | **w** |
|---|---|---|---|
| BM1 | -257.05 | 21.01 | 0.00 |
| **OU1** | **-278.06** | **0.00** | **0.42** |
| OUM$_{loc3a}$ | -276.13 | 1.93 | 0.16 |
| OUM$_{loc3b}$ | -276.53 | 1.53 | 0.20 |
| OUM$_{loc3c}$ | -274.98 | 3.08 | 0.09 |
| OUM$_{loc4}$ | -275.79 | 2.27 | 0.13 |

| **ppxw** | **fit** | **delta** | **w** |
|---|---|---|---|
| BM1 | -161.50 | 131.90 | 0.00 |
| OU1 | -288.94 | 4.46 | 0.04 |
| **OUM$_{loc3a}$** | **-293.40** | **0.00** | **0.36** |
| OUM$_{loc3b}$ | -293.32 | 0.08 | 0.35 |
| OUM$_{loc3c}$ | -291.10 | 2.30 | 0.11 |
| OUM$_{loc4}$ | -291.43 | 1.97 | 0.14 |

| **tl** | **fit** | **delta** | **w** |
|---|---|---|---|
| BM1 | -364.70 | 22.93 | 0.00 |
| OU1 | -382.77 | 4.86 | 0.03 |
| OUM$_{loc3a}$ | -387.32 | 0.31 | 0.28 |
| OUM$_{loc3b}$ | -384.09 | 3.54 | 0.06 |
| OUM$_{loc3c}$ | -387.55 | 0.09 | 0.31 |
| **OUM$_{loc4}$** | **-387.63** | **0.00** | **0.33** |

| **tmw** | **fit** | **delta** | **w** |
|---|---|---|---|
| BM1 | -309.05 | 78.44 | 0.00 |
| **OU1** | **-387.49** | **0.00** | **0.46** |
| OUM$_{loc3a}$ | -386.13 | 1.36 | 0.23 |
| OUM$_{loc3b}$ | -385.31 | 2.18 | 0.16 |
| OUM$_{loc3c}$ | -384.33 | 3.16 | 0.09 |
| OUM$_{loc4}$ | -383.24 | 4.26 | 0.05 |

| **tpw** | **fit** | **delta** | **w** |
|---|---|---|---|
| BM1 | -352.62 | 108.62 | 0.00 |
| OU1 | -449.69 | 11.56 | 0.00 |
| OUM$_{loc3a}$ | -459.33 | 1.92 | 0.18 |
| OUM$_{loc3b}$ | -459.65 | 1.60 | 0.22 |
| **OUM$_{loc3c}$** | **-461.25** | **0.00** | **0.48** |
| OUM$_{loc4}$ | -458.45 | 2.79 | 0.12 |

**Table S3: Optima and bootstrapping results for best-supported OUM model.** We bootstrapped traits with supported OUM models and excluded traits with supported OU1 (proximal phalanx width, tibia midshaft width, pedal proximal phalanx length) for the a) forelimb and b) hind limbs. Trait abbreviations are explained in Figure S1 and table S1.

**a. forelimb mean ϴ values and 95% confidence intervals for 13 traits (1 excluded) for the best supported model.**

| **sl** | **$OUM_{loc3b}$ optima** | **L95** | **U95** |
|---|---|---|---|
| ground | 0.58 | 0.56 | 0.60 |
| arb+glide | 0.59 | 0.57 | 0.61 |
| fly | 0.67 | 0.64 | 0.71 |

| **sh** | **$OUM_{loc4}$ optima** | **L95** | **U95** |
|---|---|---|---|
| ground | 0.30 | 0.28 | 0.33 |
| arb. | 0.32 | 0.28 | 0.35 |
| glide | 0.24 | 0.15 | 0.33 |
| fly | 0.51 | 0.46 | 0.57 |

| **hl** | **$OUM_{loc4}$ optima** | **L95** | **U95** |
|---|---|---|---|
| ground | 0.63 | 0.61 | 0.66 |
| arb. | 0.74 | 0.70 | 0.77 |
| glide | 0.93 | 0.83 | 1.03 |
| fly | 1.10 | 1.03 | 1.16 |

| **hdw** | **$OUM_{loc3b}$ optima** | **L95** | **U95** |
|---|---|---|---|
| ground | 0.09 | 0.06 | 0.13 |
| arb+glide | 0.15 | 0.10 | 0.20 |
| fly | 0.22 | 0.14 | 0.31 |

| **hsw** | **$OUM_{loc3b}$ optima** | **L95** | **U95** |
|---|---|---|---|
| ground | -0.40 | -0.42 | -0.38 |
| arb+glide | -0.34 | -0.37 | -0.32 |
| fly | -0.23 | -0.27 | -0.18 |

| **hpw** | **$OUM_{loc3b}$ optima** | **L95** | **U95** |
|---|---|---|---|
| ground | 0.01 | -0.01 | 0.03 |
| arb+glide | 0.01 | -0.01 | 0.03 |
| fly | 0.18 | 0.14 | 0.22 |

| **rl** | **$OUM_{loc4}$ optima** | **L95** | **U95** |
|---|---|---|---|
| ground | 0.60 | 0.57 | 0.63 |
| arb. | 0.73 | 0.68 | 0.77 |
| glide | 1.00 | 0.87 | 1.13 |
| fly | 1.33 | 1.24 | 1.41 |

| **ul** | **$OUM_{loc4}$ optima** | **L95** | **U95** |
|---|---|---|---|
| ground | 0.71 | 0.69 | 0.73 |
| arb. | 0.79 | 0.76 | 0.82 |
| glide | 0.97 | 0.89 | 1.05 |
| fly | 1.29 | 1.24 | 1.33 |

| **uol** | **$OUM_{loc3a}$ optima** | **L95** | **U95** |
|---|---|---|---|
| ground | -0.09 | -0.11 | -0.06 |
| arb. | -0.18 | -0.22 | -0.15 |

| **mcl** | **$OUM_{loc3c}$ optima** | **L95** | **U95** |
|---|---|---|---|
| ground+arb | 0.04 | -0.08 | 0.15 |
| glide | 0.24 | -0.31 | 0.78 |

| | | | |
|---|---|---|---|
| glide+fly | -0.28 | -0.33 | -0.23 |

| **mcw** | **OUM$_{loc3b}$ optima** | **L95** | **U95** |
|---|---|---|---|
| ground | -0.71 | -0.75 | -0.68 |
| arb+glide | -0.77 | -0.83 | -0.71 |
| fly | -0.59 | -0.69 | -0.49 |

| **ipl** | **OUM$_{loc4}$ optima** | **L95** | **U95** |
|---|---|---|---|
| ground | -0.32 | -0.37 | -0.28 |
| arb. | -0.13 | -0.21 | -0.06 |
| glide | 0.17 | -0.05 | 0.38 |
| **fly** | 1.06 | 0.85 | 1.25 |

| | | | |
|---|---|---|---|
| fly | 2.41 | 1.50 | 3.68 |

| **ppl** | **OUM$_{loc4}$ optima** | **L95** | **U95** |
|---|---|---|---|
| ground | -0.22 | -0.28 | -0.17 |
| arb. | 0.06 | -0.05 | 0.15 |
| glide | 0.28 | 0.01 | 0.52 |
| **fly** | 1.14 | 0.83 | 1.39 |

### b. hind limb mean ϴ values and 95% confidence intervals for 13 traits (2 excluded) for the best supported model.

| **fbl** | **OUM$_{loc4}$ optima** | **L95** | **U95** |
|---|---|---|---|
| ground | 0.76 | 0.72 | 0.80 |
| arb. | 0.85 | 0.78 | 0.91 |
| glide | 1.05 | 0.85 | 1.23 |
| fly | 0.72 | 0.61 | 0.82 |

| **fl** | **OUM$_{loc4}$ optima** | **L95** | **U95** |
|---|---|---|---|
| ground | 0.72 | 0.69 | 0.75 |
| arb. | 0.85 | 0.79 | 0.89 |
| glide | 1.01 | 0.87 | 1.14 |
| fly | 0.86 | 0.78 | 0.94 |

| **il** | **OUM$_{loc3b}$ optima** | **L95** | **U95** |
|---|---|---|---|
| ground | 0.54 | 0.51 | 0.56 |
| arb+glide | 0.59 | 0.56 | 0.62 |
| fly | 0.45 | 0.39 | 0.51 |

| **fdw** | **OUM$_{loc3c}$ optima** | **L95** | **U95** |
|---|---|---|---|
| ground+arb | 0.06 | 0.05 | 0.07 |
| glide | 0.03 | -0.02 | 0.08 |
| fly | -0.05 | -0.09 | -0.01 |

| **fsw** | **OUM$_{loc3a}$ optima** | **L95** | **U95** |
|---|---|---|---|
| ground | -0.29 | -0.31 | -0.27 |
| arb. | -0.29 | -0.31 | -0.26 |
| glide+fly | -0.36 | -0.40 | -0.33 |

| **ipxl** | **OUM$_{loc3b}$ optima** | **L95** | **U95** |
|---|---|---|---|
| ground | -0.20 | -0.24 | -0.17 |
| arb+glide | -0.12 | -0.16 | -0.08 |
| fly | -0.05 | -0.13 | 0.03 |

| **mtl** | **OUM$_{loc3b}$ optima** | **L95** | **U95** |
|---|---|---|---|

| isl | $OUM_{loc3b}$ optima | L95 | U95 |
|---|---|---|---|
| ground | 0.37 | 0.35 | 0.40 |
| arb+glide | 0.31 | 0.28 | 0.35 |
| fly | 0.13 | 0.07 | 0.20 |

| mtw | $OUM_{loc3a}$ optima | L95 | U95 |
|---|---|---|---|
| ground | -0.64 | -0.67 | -0.61 |
| arb. | -0.71 | -0.75 | -0.67 |
| glide+fly | -0.79 | -0.84 | -0.73 |

| ppxw | $OUM_{loc3a}$ optima | L95 | U95 |
|---|---|---|---|
| ground | -0.69 | -0.72 | -0.67 |
| arb. | -0.73 | -0.77 | -0.70 |
| glide+fly | -0.77 | -0.82 | -0.72 |

| tpw | $OUM_{loc3c}$ optima | L95 | U95 |
|---|---|---|---|
| ground+arb | 0.05 | 0.03 | 0.06 |
| glide | 0.01 | -0.04 | 0.06 |
| fly | -0.04 | -0.08 | 0.00 |

| | | | |
|---|---|---|---|
| ground | 0.37 | 0.27 | 0.48 |
| arb+glide | 0.17 | 0.05 | 0.32 |
| fly | -0.03 | -0.28 | 0.25 |

| pel | $OUM_{loc3c}$ optima | L95 | U95 |
|---|---|---|---|
| ground+arb | 0.77 | 0.76 | 0.79 |
| glide | 0.79 | 0.71 | 0.87 |
| fly | 0.62 | 0.57 | 0.67 |

| tl | $OUM_{loc4}$ optima | L95 | U95 |
|---|---|---|---|
| ground | 0.79 | 0.74 | 0.83 |
| arb. | 0.86 | 0.78 | 0.93 |
| glide | 1.09 | 0.86 | 1.28 |
| fly | 0.89 | 0.76 | 1.00 |

**Table S4: Statistical results for model fitting analyses on the extant+extinct dataset.** Comparisons of the best-fitting OUM models (bolded) using AICc scores to asses fit and AICcW weights to asses relative model support for a) 14 forelimb traits and b) 15 hind limb traits. We fit all models using functions in the *OUwie R* package. Trait abbreviations are explained in table S1.

a. Forelimb

| **hdw** | **fit** | **delta** | **w** |
|---|---|---|---|
| BM1 | -414.31 | 36.66 | 0 |
| OU1 | -448.77 | 2.2 | 0.14 |
| OUM$_{loc3a}$ | -448.46 | 2.51 | 0.12 |
| **OUM$_{loc3b}$** | **-450.97** | **0** | **0.41** |
| OUM$_{loc3c}$ | -448.86 | 2.11 | 0.14 |
| OUM$_{loc4}$ | -449.54 | 1.43 | 0.2 |

| **hl** | **fit** | **delta** | **w** |
|---|---|---|---|
| BM1 | -414.17 | 87.35 | 0 |
| OU1 | -435.55 | 65.97 | 0 |
| OUM$_{loc3a}$ | -489.84 | 11.68 | 0 |
| OUM$_{loc3b}$ | -489.51 | 12.01 | 0 |
| OUM$_{loc3c}$ | -490.29 | 11.23 | 0 |
| **OUM$_{loc4}$** | **-501.52** | **0** | **0.99** |

| **hpw** | **fit** | **delta** | **w** |
|---|---|---|---|
| BM1 | -469.58 | 89.9 | 0 |
| OU1 | -523.52 | 35.96 | 0 |
| **OUM$_{loc3a}$** | **-559.48** | **0** | **0.35** |
| OUM$_{loc3b}$ | -559 | 0.48 | 0.27 |
| OUM$_{loc3c}$ | -558.82 | 0.66 | 0.25 |
| OUM$_{loc4}$ | -557.53 | 1.95 | 0.13 |

| **hsw** | **fit** | **delta** | **w** |
|---|---|---|---|
| BM1 | -364 | 72.66 | 0 |
| OU1 | -416.5 | 20.17 | 0 |
| OUM$_{loc3a}$ | -430.17 | 6.5 | 0.03 |
| **OUM$_{loc3b}$** | **-436.67** | **0** | **0.66** |
| OUM$_{loc3c}$ | -429.92 | 6.74 | 0.02 |
| OUM$_{loc4}$ | -435.04 | 1.63 | 0.29 |

| **ipl** | **fit** | **delta** | **w** |
|---|---|---|---|
| BM1 | -182.49 | 106.48 | 0 |
| OU1 | -191.73 | 97.24 | 0 |
| OUM$_{loc3a}$ | -280.35 | 8.62 | 0.01 |
| OUM$_{loc3b}$ | -283.97 | 5 | 0.07 |
| OUM$_{loc3c}$ | -281.14 | 7.83 | 0.02 |
| **OUM$_{loc4}$** | **-288.97** | **0** | **0.9** |

| **mcl** | **fit** | **delta** | **w** |
|---|---|---|---|
| BM1 | -292.42 | 53.72 | 0 |
| OU1 | -295.49 | 50.65 | 0 |
| OUM$_{loc3a}$ | -340.94 | 5.2 | 0.04 |
| **OUM$_{loc3b}$** | **-346.14** | **0** | **0.58** |
| OUM$_{loc3c}$ | -345.19 | 0.95 | 0.36 |
| OUM$_{loc4}$ | -338.44 | 7.7 | 0.01 |

| **mcw** | **fit** | **delta** | **w** |
|---|---|---|---|
| BM1 | -319.17 | 40.19 | 0 |
| OU1 | -356.63 | 2.74 | 0.1 |
| OUM$_{loc3a}$ | -357.84 | 1.52 | 0.19 |
| **OUM$_{loc3b}$** | **-359.36** | **0** | **0.4** |
| OUM$_{loc3c}$ | -357.69 | 1.67 | 0.17 |
| OUM$_{loc4}$ | -357.2 | 2.16 | 0.14 |

| **ppl** | **fit** | **delta** | **w** |
|---|---|---|---|
| BM1 | -299.33 | 74.17 | 0 |
| OU1 | -300.18 | 73.32 | 0 |
| OUM$_{loc3a}$ | -351.81 | 21.69 | 0 |
| OUM$_{loc3b}$ | -372.28 | 1.22 | 0.35 |
| OUM$_{loc3c}$ | -351.11 | 22.39 | 0 |
| **OUM$_{loc4}$** | **-373.5** | **0** | **0.65** |

| **ppw** | **fit** | **delta** | **w** |
|---|---|---|---|
| BM1 | -305.14 | 47.54 | 0 |
| **OU1** | **-352.68** | **0** | **0.55** |
| OUM$_{loc3a}$ | -348.69 | 3.99 | 0.08 |
| OUM$_{loc3b}$ | -350.86 | 1.82 | 0.22 |
| OUM$_{loc3c}$ | -349.03 | 3.65 | 0.09 |
| OUM$_{loc4}$ | -348.13 | 4.55 | 0.06 |

| **rl** | fit | delta | w |
|---|---|---|---|
| BM1 | -357.08 | 83.62 | 0 |
| OU1 | -365.86 | 74.84 | 0 |
| OUM$_{loc3a}$ | -427.88 | 12.82 | 0 |
| OUM$_{loc3b}$ | -427 | 13.7 | 0 |

| sh | fit | delta | w |
|---|---|---|---|
| BM1 | -315.44 | 89.03 | 0 |
| OU1 | -376.83 | 27.64 | 0 |
| **OUM$_{loc3a}$** | **-404.47** | **0** | **0.42** |
| OUM$_{loc3b}$ | -401.68 | 2.79 | 0.1 |

| sl | fit | delta | w |
|---|---|---|---|
| BM1 | -393.8 | 133.47 | 0 |
| OU1 | -506.44 | 20.83 | 0 |
| OUM$_{loc3a}$ | -526.78 | 0.49 | 0.26 |
| **OUM$_{loc3b}$** | **-527.27** | **0** | **0.33** |

| | | | |
|---|---|---|---|
| OUM$_{loc3c}$ | -429.04 | 11.66 | 0 |
| **OUM$_{loc4}$** | **-440.7** | **0** | **0.99** |

| | | | |
|---|---|---|---|
| OUM$_{loc3c}$ | -403.43 | 1.04 | 0.25 |
| OUM$_{loc4}$ | -403.29 | 1.19 | 0.23 |

| | | | |
|---|---|---|---|
| OUM$_{loc3c}$ | -527.05 | 0.22 | 0.3 |
| OUM$_{loc4}$ | -524.93 | 2.34 | 0.1 |

| **ul** | fit | delta | w |
|---|---|---|---|
| BM1 | -401.44 | 124.63 | 0 |
| OU1 | -409.77 | 116.3 | 0 |
| OUM$_{loc3a}$ | -509.25 | 16.82 | 0 |
| OUM$_{loc3b}$ | -498.37 | 27.7 | 0 |
| OUM$_{loc3c}$ | -502.25 | 23.82 | 0 |
| **OUM$_{loc4}$** | **-526.07** | **0** | **1** |

| **uol** | **fit** | **delta** | **w** |
|---|---|---|---|
| BM1 | -264.02 | 82.47 | 0 |
| OU1 | -314.8 | 31.69 | 0 |
| OUM$_{loc3a}$ | -334.21 | 12.28 | 0 |
| OUM$_{loc3b}$ | -345.14 | 1.35 | 0.34 |
| OUM$_{loc3c}$ | -333.88 | 12.6 | 0 |
| **OUM$_{loc4}$** | **-346.49** | **0** | **0.66** |

b. Hind limb

| **fbl** | **fit** | **delta** | **w** |
|---|---|---|---|
| BM1 | -336.31 | 24.84 | 0 |
| OU1 | -356.4 | 4.76 | 0.05 |
| OUM$_{loc3a}$ | -355.13 | 6.03 | 0.03 |
| OUM$_{loc3b}$ | -357.8 | 3.35 | 0.11 |
| OUM$_{loc3c}$ | -359.53 | 1.62 | 0.25 |
| **OUM$_{loc4}$** | **-361.15** | **0** | **0.56** |

| **fdw** | **fit** | **delta** | **w** |
|---|---|---|---|
| BM1 | -464.14 | 95.9 | 0 |
| OU1 | -536.38 | 23.66 | 0 |
| OUM$_{loc3a}$ | -556.29 | 3.75 | 0.06 |
| OUM$_{loc3b}$ | -559.52 | 0.52 | 0.29 |
| **OUM$_{loc3c}$** | **-560.04** | **0** | **0.37** |
| OUM$_{loc4}$ | -559.46 | 0.58 | 0.28 |

| **fl** | **fit** | **delta** | **w** |
|---|---|---|---|
| BM1 | -399.43 | 34.99 | 0 |
| OU1 | -417.78 | 16.64 | 0 |
| OUM$_{loc3a}$ | -431.38 | 3.03 | 0.17 |
| OUM$_{loc3b}$ | -429.49 | 4.93 | 0.06 |
| OUM$_{loc3c}$ | -423.86 | 10.56 | 0 |
| **OUM$_{loc4}$** | **-434.41** | **0** | **0.76** |

| **fsw** | **fit** | **delta** | **w** |
|---|---|---|---|
| BM1 | -419.08 | 80.97 | 0 |
| OU1 | -492.23 | 7.83 | 0.01 |
| **OUM$_{loc3a}$** | **-500.05** | **0** | **0.42** |
| OUM$_{loc3b}$ | -496.49 | 3.57 | 0.07 |
| OUM$_{loc3c}$ | -499.82 | 0.24 | 0.37 |
| OUM$_{loc4}$ | -497.82 | 2.23 | 0.14 |

| **il** | **fit** | **delta** | **w** |
|---|---|---|---|
| BM1 | -428.09 | 50.06 | 0 |
| OU1 | -471.26 | 6.89 | 0.02 |
| OUM$_{loc3a}$ | -473.08 | 5.06 | 0.04 |
| **OUM$_{loc3b}$** | **-478.15** | **0** | **0.54** |
| OUM$_{loc3c}$ | -474.07 | 4.08 | 0.07 |
| OUM$_{loc4}$ | -477.21 | 0.94 | 0.34 |

| **ipxl** | **fit** | **delta** | **w** |
|---|---|---|---|
| BM1 | -99.84 | 97.13 | 0 |
| OU1 | -186.3 | 10.67 | 0 |
| OUM$_{loc3a}$ | -195.94 | 1.03 | 0.28 |
| **OUM$_{loc3b}$** | **-196.97** | **0** | **0.47** |
| OUM$_{loc3c}$ | -191.71 | 5.26 | 0.03 |
| OUM$_{loc4}$ | -195.29 | 1.68 | 0.2 |

| **isl** | **fit** | **delta** | **w** |
|---|---|---|---|
| BM1 | -362.54 | 59.58 | 0 |
| OU1 | -393.96 | 28.16 | 0 |
| OUM$_{loc3a}$ | -416.29 | 5.82 | 0.04 |
| **OUM$_{loc3b}$** | **-422.12** | **0** | **0.76** |
| OUM$_{loc3c}$ | -415.33 | 6.79 | 0.03 |
| OUM$_{loc4}$ | -419.17 | 2.94 | 0.17 |

| **mtl** | **fit** | **delta** | **w** |
|---|---|---|---|
| BM1 | -234.47 | 15.71 | 0 |
| OU1 | -244.72 | 5.46 | 0.03 |
| OUM$_{loc3a}$ | -249.61 | 0.57 | 0.35 |
| **OUM$_{loc3b}$** | **-250.18** | **0** | **0.46** |
| OUM$_{loc3c}$ | -244.74 | 5.45 | 0.03 |
| OUM$_{loc4}$ | -247.6 | 2.58 | 0.13 |

| **pel** | **fit** | **delta** | **w** |
|---|---|---|---|
| BM1 | -446.28 | 59.11 | 0 |
| OU1 | -489.28 | 16.11 | 0 |
| OUM$_{loc3a}$ | -498.44 | 6.94 | 0.01 |
| **OUM$_{loc3b}$** | **-505.38** | **0** | **0.43** |
| OUM$_{loc3c}$ | -505.19 | 0.2 | 0.39 |
| OUM$_{loc4}$ | -503.58 | 1.81 | 0.17 |

| **tl** | **fit** | **delta** | **w** |
|---|---|---|---|
| BM1 | -370.43 | 22.64 | 0 |

| **tmw** | **fit** | **delta** | **w** |
|---|---|---|---|
| BM1 | -300.03 | 85.01 | 0 |

| **tpw** | **fit** | **delta** | **w** |
|---|---|---|---|
| BM1 | -358.46 | 107.88 | 0 |

| | | | |
|---|---|---|---|
| OU1 | -388.18 | 4.9 | 0.04 |
| $OUM_{loc3a}$ | -391.47 | 1.6 | 0.19 |
| $OUM_{loc3b}$ | -389.18 | 3.9 | 0.06 |
| $OUM_{loc3c}$ | -392.24 | 0.83 | 0.28 |
| **$OUM_{loc4}$** | **-393.07** | **0** | **0.43** |

| | | | |
|---|---|---|---|
| **OU1** | **-385.04** | **0** | **0.44** |
| $OUM_{loc3a}$ | -383.38 | 1.66 | 0.19 |
| $OUM_{loc3b}$ | -383.46 | 1.58 | 0.2 |
| $OUM_{loc3c}$ | -382 | 3.04 | 0.1 |
| $OUM_{loc4}$ | -381.33 | 3.71 | 0.07 |

| | | | |
|---|---|---|---|
| OU1 | -450.83 | 15.52 | 0 |
| $OUM_{loc3a}$ | -464.22 | 2.13 | 0.16 |
| $OUM_{loc3b}$ | -464.86 | 1.49 | 0.22 |
| **$OUM_{loc3c}$** | **-466.35** | **0** | **0.46** |
| $OUM_{loc4}$ | -464.32 | 2.03 | 0.17 |

| **ppxw** | **fit** | **delta** | **w** |
|---|---|---|---|
| BM1 | -153.48 | 136.46 | 0 |
| OU1 | -283.09 | 6.86 | 0.01 |
| **$OUM_{loc3a}$** | **-289.94** | **0** | **0.39** |
| $OUM_{loc3b}$ | -289.66 | 0.28 | 0.34 |
| $OUM_{loc3c}$ | -287.42 | 2.52 | 0.11 |
| $OUM_{loc4}$ | -288.12 | 1.82 | 0.16 |

| **ppxl** | **fit** | **delta** | **w** |
|---|---|---|---|
| BM1 | -253.68 | 22.28 | 0 |
| **OU1** | **-275.97** | **0** | **0.44** |
| $OUM_{loc3a}$ | -274.31 | 1.66 | 0.19 |
| $OUM_{loc3b}$ | -274.44 | 1.53 | 0.2 |
| $OUM_{loc3c}$ | -272.48 | 3.48 | 0.08 |
| $OUM_{loc4}$ | -272.94 | 3.02 | 0.1 |

| **mtw** | **fit** | **delta** | **w** |
|---|---|---|---|
| BM1 | -72.59 | 160.39 | 0 |
| OU1 | -212.33 | 20.65 | 0 |
| **$OUM_{loc3a}$** | **-232.98** | **0** | **0.58** |
| $OUM_{loc3b}$ | -230.79 | 2.19 | 0.19 |
| $OUM_{loc3c}$ | -226.85 | 6.13 | 0.03 |
| $OUM_{loc4}$ | -230.89 | 2.09 | 0.2 |

**Table S5: Optima values for the best-fitting OUM model results for extant+extinct dataset.** Bootstrapping did not perform well on this dataset, so here we include only the model optima values with upper and lower 95% confidence intervals for a) forelimb traits and b) hind limb traits. We excluded traits only supported by an OU1 model (proximal phalanx width and pedal proximal phalanx length). Trait abbreviations are explained in Figure S1 and table S1.

### a. forelimb

| **sl** | **OUMloc3b optima** | **L95** | **U95** |
|---|---|---|---|
| **ground** | 0.58 | 0.58 | 0.58 |
| **arb+glide** | 0.59 | 0.59 | 0.59 |
| **fly** | 0.68 | 0.68 | 0.68 |

| **sh** | **OUMloc3a optima** | **L95** | **U95** |
|---|---|---|---|
| ground | 0.31 | 0.31 | 0.31 |
| arb. | 0.24 | 0.23 | 0.26 |
| glide+fly | 0.50 | 0.50 | 0.51 |

| **uol** | **OUMloc4 optima** | **L95** | **U95** |
|---|---|---|---|
| **ground** | -0.09 | -0.09 | -0.08 |
| **arb.** | -0.18 | -0.19 | -0.17 |
| **glide** | -0.27 | -0.31 | -0.22 |
| **fly** | -0.28 | -0.28 | -0.27 |

| **hdw** | **OUMloc3b optima** | **L95** | **U95** |
|---|---|---|---|
| ground | 0.09 | 0.07 | 0.11 |
| arb. & glide | 0.15 | 0.12 | 0.17 |
| fly | 0.21 | 0.21 | 0.21 |

| **hsw** | **OUMloc3b optima** | **L95** | **U95** |
|---|---|---|---|
| **ground** | -0.40 | -0.41 | -0.40 |
| **arb. & glide** | -0.35 | -0.35 | -0.33 |
| **fly** | -0.21 | -0.21 | -0.21 |

| **hpw** | **OUMloc3a optima** | **L95** | **U95** |
|---|---|---|---|
| ground | 0.01 | 0.01 | 0.01 |
| arb. | 0.00 | -0.01 | 0.00 |
| glide+fly | 0.19 | 0.19 | 0.19 |

| **rl** | **OUMloc4 optima** | **L95** | **U95** |
|---|---|---|---|
| **ground** | 0.61 | 0.60 | 0.62 |

| **ul** | **OUMloc4 optima** | **L95** | **U95** |
|---|---|---|---|
| ground | 0.71 | 0.71 | 0.72 |

| | | | |
|---|---|---|---|
| **arb.** | 0.72 | 0.70 | 0.73 |
| **glide** | 0.93 | 0.87 | 1.01 |
| **fly** | 1.29 | 1.29 | 1.31 |

| | | | |
|---|---|---|---|
| arb. | 0.79 | 0.78 | 0.80 |
| glide | 0.94 | 0.91 | 0.97 |
| fly | 1.29 | 1.29 | 1.29 |

| **mcl** | **OUMloc3b optima** | **L95** | **U95** |
|---|---|---|---|
| **ground** | -0.04 | -0.10 | 0.01 |
| **arb. & glide** | 0.03 | -0.05 | 0.08 |
| **fly** | 1.44 | 1.37 | 1.48 |

| **mcw** | **OUMloc3b optima** | **L95** | **U95** |
|---|---|---|---|
| ground | -0.72 | -0.73 | -0.70 |
| arb. & glide | -0.77 | -0.80 | -0.74 |
| fly | -0.60 | -0.61 | -0.60 |

| **hl** | **OUMloc4 optima** | **L95** | **U95** |
|---|---|---|---|
| **ground** | 0.64 | 0.63 | 0.64 |
| **arb.** | 0.73 | 0.72 | 0.75 |
| **glide** | 0.93 | 0.86 | 0.99 |
| **fly** | 1.09 | 1.09 | 1.10 |

| **ppl** | **OUMloc4 optima** | **L95** | **U95** |
|---|---|---|---|
| ground | -0.23 | -0.28 | -0.19 |
| arb. | 0.04 | -0.02 | 0.10 |
| glide | 0.24 | 0.15 | 0.31 |
| fly | 0.92 | 0.89 | 0.95 |

| **ipl** | **OUMloc4 optima** | **L95** | **U95** |
|---|---|---|---|
| **ground** | -0.32 | -0.34 | -0.30 |
| **arb.** | -0.15 | -0.19 | -0.12 |
| **glide** | 0.13 | 0.02 | 0.22 |
| **fly** | 0.95 | 0.93 | 0.97 |

### b. hind limb

| **pel** | **OUMloc4b optima** | **L95** | **U95** |
|---|---|---|---|
| **ground** | 0.77 | 0.77 | 0.78 |
| **arb+glide** | 0.78 | 0.77 | 0.79 |
| **fly** | 0.63 | 0.63 | 0.63 |

| **il** | **OUMloc3b optima** | **L95** | **U95** |
|---|---|---|---|
| ground | 0.54 | 0.53 | 0.55 |
| arb+glide | 0.59 | 0.57 | 0.60 |
| fly | 0.46 | 0.46 | 0.46 |

| **isl** | **OUMloc3b optima** | **L95** | **U95** |
|---|---|---|---|
| **ground** | 0.38 | 0.37 | 0.38 |
| **arb+glide** | 0.31 | 0.29 | 0.32 |
| **fly** | 0.13 | 0.13 | 0.13 |

| **fsw** | **OUMloc3a optima** | **L95** | **U95** |
|---|---|---|---|
| ground | -0.29 | -0.29 | -0.29 |
| arb. | -0.36 | -0.37 | -0.35 |
| glide+fly | -0.35 | -0.35 | -0.35 |

| **fl** | **OUMloc4 optima** | **L95** | **U95** |
|---|---|---|---|
| **ground** | 0.72 | 0.71 | 0.73 |
| **arb.** | 0.85 | 0.81 | 0.88 |
| **glide** | 1.03 | 0.98 | 1.12 |
| **fly** | 0.86 | 0.85 | 0.86 |

| **tl** | **OUMloc4 optima** | **L95** | **U95** |
|---|---|---|---|
| ground | 0.79 | 0.77 | 0.80 |
| arb. | 0.86 | 0.82 | 0.90 |
| glide | 1.12 | 1.04 | 1.23 |
| fly | 0.88 | 0.87 | 0.88 |

| **fbl** | **OUMloc4 optima** | **L95** | **U95** |
|---|---|---|---|

| **tpw** | **OUMloc3c optima** | **L95** | **U95** |
|---|---|---|---|

| | | | |
|---|---|---|---|
| **ground** | 0.76 | 0.74 | 0.77 |
| **arb.** | 0.84 | 0.80 | 0.88 |
| **glide** | 1.10 | 1.03 | 1.21 |
| **fly** | 0.71 | 0.70 | 0.72 |

| | | | |
|---|---|---|---|
| ground+arb | 0.05 | 0.05 | 0.05 |
| glide | 0.01 | 0.00 | 0.02 |
| fly | -0.05 | -0.05 | -0.05 |

| **mtl** | **OUMloc3b optima** | **L95** | **U95** |
|---|---|---|---|
| **ground** | 0.39 | 0.36 | 0.43 |
| **arb+glide** | 0.19 | 0.12 | 0.25 |
| **fly** | 0.09 | 0.08 | 0.10 |

| **mtw** | **OUMloc3a optima** | **L95** | **U95** |
|---|---|---|---|
| ground | -0.64 | -0.65 | -0.64 |
| arb. | -0.71 | -0.72 | -0.70 |
| glide+fly | -0.80 | -0.80 | -0.79 |

| **ipxl** | **optima** | **L95** | **U95** |
|---|---|---|---|
| **ground** | -0.20 | -0.21 | -0.20 |
| **arb+glide** | -0.12 | -0.13 | -0.12 |
| **fly** | -0.05 | -0.05 | -0.05 |

| **ppxw** | **OUMloc3a optima** | **L95** | **U95** |
|---|---|---|---|
| ground | -0.70 | -0.70 | -0.69 |
| arb. | -0.73 | -0.74 | -0.73 |
| glide+fly | -0.79 | -0.79 | -0.78 |

| **fdw** | **OUMloc3c optima** | **L95** | **U95** |
|---|---|---|---|
| **ground+arb** | 0.06 | 0.06 | 0.06 |
| **glide** | 0.03 | 0.01 | 0.04 |
| **fly** | -0.05 | -0.05 | -0.05 |

**Table S6.** Fits of multivariate evolutionary models to morphological data of extant+extinct species. A) MvMORPH models were fit to PC1–PC3 scores of a principal component analysis PCA of 29 skeletal traits. B) Optima were extracted from the model, but were physically impossible for the forelimb models so cannot be properly analyzed.

a)

| **forelimb** | **fit** | **delta** | **w** |
|---|---|---|---|
| mvBM1 | -514.32 | 239.51 | 0.00 |
| mvOU1 | -561.39 | 192.44 | 0.00 |
| mvOUM$_{loc3a}$ | -722.22 | 31.60 | 0.00 |
| mvOUM$_{loc3b}$ | -730.87 | 22.95 | 0.00 |
| mvOUM$_{loc3c}$ | -712.38 | 41.45 | 0.00 |
| **mvOUM$_{loc4}$** | **-753.83** | **0.00** | **1.00** |

| **hind limb** | **fit** | **delta** | **w** |
|---|---|---|---|
| mvBM1 | -241.16 | 212.95 | 0.00 |
| mvOU1 | -400.68 | 53.43 | 0.00 |
| mvOUM$_{loc3a}$ | -450.88 | 3.24 | 0.16 |
| mvOUM$_{loc3b}$ | -447.86 | 6.25 | 0.04 |
| mvOUM$_{loc3c}$ | -435.10 | 19.01 | 0.00 |
| **mvOUM$_{loc4}$** | **-454.11** | **0.00** | **0.80** |

b) PC1 and PC2 optima are formatted as points (PC1, PC2) that could be superimposed on a PCA.

| best supported model | ground-dweller optimum | arborealist optimum | glider optimum | bat optimum |
|---|---|---|---|---|
| forelimb mvMORPH$_{loc4}$ PC1 value | -118.91 | -100.66 | -55.78 | -6.85 |
| forelimb mvMORPH$_{loc4}$ PC2 value | -256.49 | -214.10 | -107.61 | -24.24 |
| hind limb mvMORPH$_{loc4}$ PC1 value | 0.03 | 0.01 | -0.01 | -0.20 |

| | | | | |
|---|---|---|---|---|
| hind limb $\text{mvMORPH}_{loc4}$ PC2 value | 0.03 | 0.00 | -0.02 | 0.20 |

**Table S7:** Summary of the best-fitting univariate OUwie models with interpretation of the optima (θ) in the context of our initial predictions (see Methods) for the a) forelimb and b) hind limb.

a) Forelimb

| Forelimb trait | Extant | θ interpretation | Supports hypothesis? | Extant+extinct |
|---|---|---|---|---|
| Scapula length | 3b | ground~arb+glide < bat | no | 3b |
| Scapula height | 4 | ground~arb~glide < bat | no | 3a |
| Humerus length | 4 | ground<arb<glide<bat | yes | 4 |
| Radius length | 4 | ground<arb<glide<bat | yes | 4 |
| Ulna length | 4 | ground<arb<glide<fly | yes | 4 |
| Ulna olecranon length | 3a | ground>arb>glide+fly | yes | 4 |
| Metacarpal length | 3c | ground+arb<glide<bat | yes | 3b |
| Prox. Phalanx length | 4 | ground<arb<glide<bat | yes | 4 |
| Intermed. Phalax length | 4 | ground<arb<glide<bat | yes | 4 |
| Humerus proximal width | 3b | ground~arb+glide<bat | no | 3a |
| Humerus shaft width | 3b | ground<arb+glide<bat | yes | 3b |
| Humerus distal width | 3b | ground<arb+glide<bat | yes | 3b |
| Metacarpal width | 3b | ground>arb+glide<bat | no | 3b |
| Prox. Phalanx width | OU1 | -- | no | OU1 |

b) Hind limb

| Hind limb trait | Extant | θ interpretation | Supports hypothesis? | Extant+extinct |
|---|---|---|---|---|
| Pelvis length | 3c | ground+arb~glide>bat | no | 3b |
| Ilium length | 3b | ground~arb+glide>fly | no | 3b |
| Ischium length | 3b | ground~arb+glide>fly | no | 3b |
| Femur length | 4 | ground<arb<glide>bat | no | 4 |
| Tibia length | 4 | ground<arb<glide>bat | no | 4 |
| Fibula length | 4 | ground~arb<glide>fly | no | 4 |
| Metatarsal length | 3b | ground>arb+glide>bat | yes | 3b |
| Pedal prox. Phalanx length | OU1 | -- | no | OU1 |
| Intermed. Pedal phalanx length | 3b | ground<arb+glide<bat | yes | 3b |
| Femur distal width | 3c | ground+arb>glide>bat | yes | 3c |
| Femur shaft width | 3a | ground~arb>glide+bat | yes | 3a |
| Tibia midshaft width | OU1 | -- | no | OU1 |
| Tibia proximal width | 3c | ground+arb>glide>bat | yes | 3c |
| Metatarsal width | 3a | ground>arb>glide+bat | yes | 3a |
| Pedal prox. Phalanx width | 3a | ground>arb>glide+bat | yes | 3a |
